## Supplementary figures and images for "GnRH pulse generator activity in mouse models of polycystic ovary syndrome"

### Suppl. Fig.1

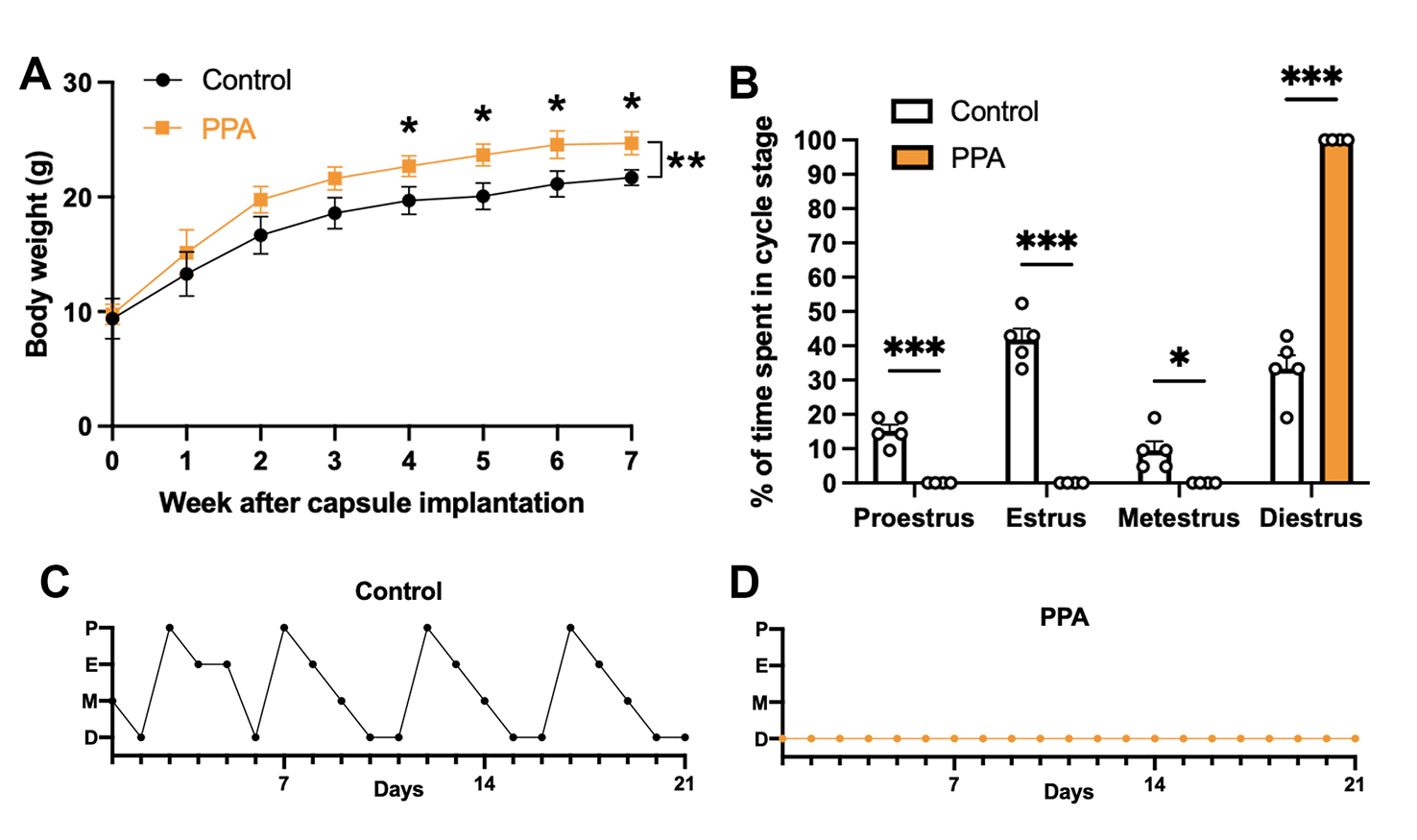

### Suppl. Fig.2

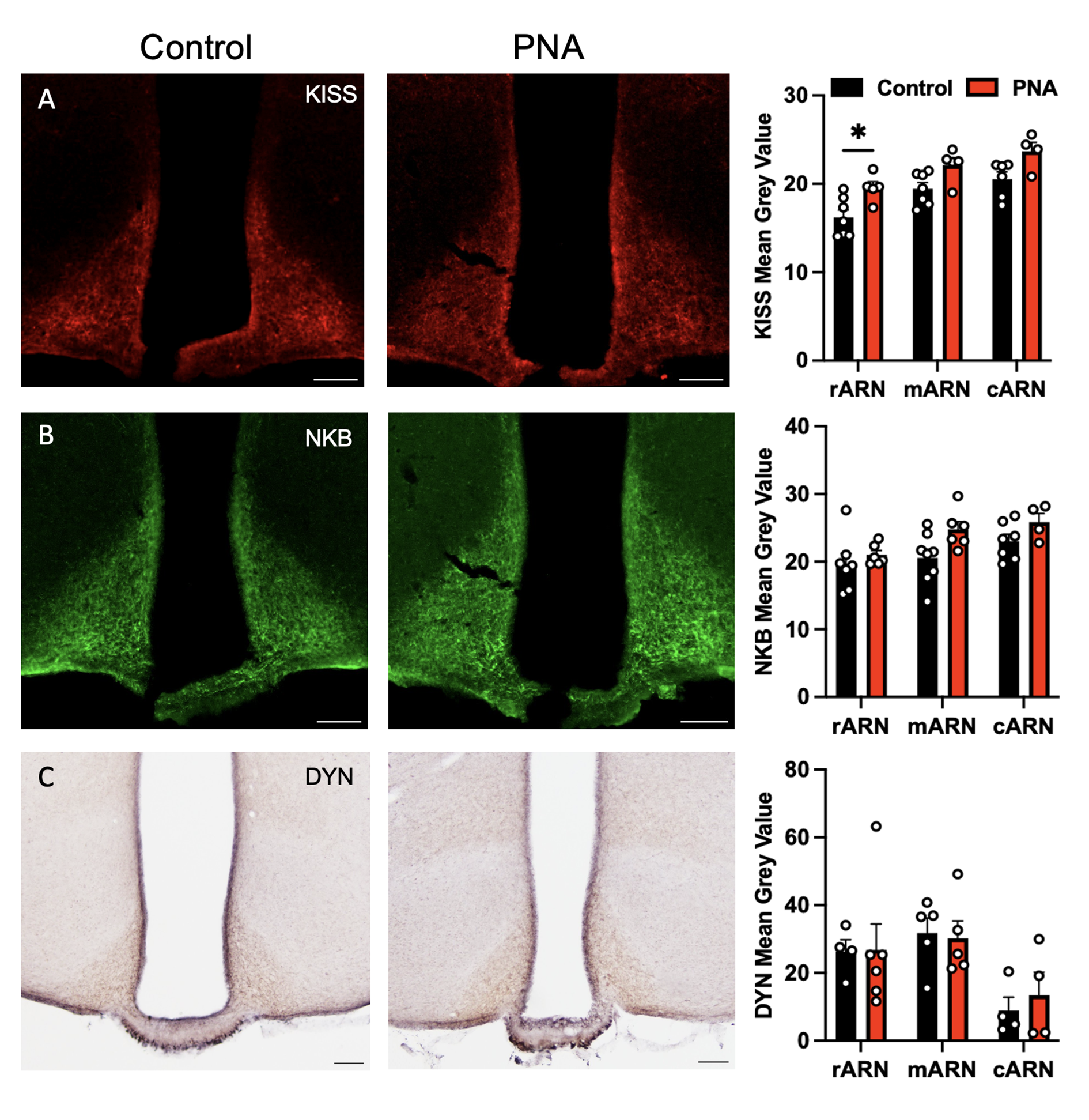
